# Transglutaminase-2 facilitates extracellular vesicle-mediated establishment of the metastatic niche

**DOI:** 10.1101/2019.12.16.875948

**Authors:** Aparna Shinde, Juan Sebastian Paez, Sarah Libring, Kelsey Hopkins, Luis Solorio, Michael K. Wendt

## Abstract

The ability of breast cancer cells to interconvert between epithelial and mesenchymal states contributes to their metastatic potential. As opposed to cell autonomous effects, the impact of epithelial-mesenchymal plasticity (EMP) on primary and metastatic tumor microenvironments remains poorly characterized. Herein we utilize global gene expression analyses to characterize a metastatic model of EMP as compared to their non-metastatic counterparts. Using this approach we demonstrate that upregulation of the extracellular matrix crosslinking enzyme tissue transglutaminase-2 (TGM2) is part of novel gene signature that only emerges in metastatic cells that have undergone induction and reversion of epithelial-mesenchymal transition (EMT). Consistent with our model system patient survival is diminished when primary tumors demonstrate enhanced levels of TGM2 in conjunction with its substrate, fibronectin. Targeted depletion of TGM2 inhibits metastasis, while overexpression of TGM2 is sufficient to enhance this process. In addition to being present within cells, we demonstrate a robust increase in the amount of TGM2 and crosslinked fibronectin present within extracellular vesicle (EV) fractions derived from metastatic breast cancer cells. Confocal microscopy of these EVs suggests that FN becomes fibrillated on their surface via a TGM2 and Tesin1-dependent process. Upon *in vivo* administration, the ability of tumor-derived EVs to induce metastatic niche formation and enhance subsequent pulmonary tumor growth requires the presence and activity of TGM2. Finally, we develop a novel 3D model of the metastatic niche to demonstrate that education of pulmonary fibroblasts via pretreatment with tumor-derived EVs promotes subsequent growth of breast cancer cells in a TGM2-dependent fashion. Overall, our studies illustrate a novel mechanism through which EMP contributes to metastatic niche development and distant metastasis via tumor-derived EVs containing abberent levels of TGM2 and fibular FN.

## Introduction

Metastatic progression is the major driver of lethality in breast cancer ^1^. Overt manifestation of macroscopic metastases is the culminating event in the multistep process of disease progression. However, recent efforts from our laboratory and others clearly indicate that molecular events take place very early in disease progression which can influence the success and failure of disseminated cells to proliferate in secondary tissues and establish metastatic disease ^2^. Among these, the molecular and phenotypic aspects of epithelial-mesenchymal transition (EMT) play a key role in maximizing the metastatic potential of mammary tumors through several mechanisms ^3^. Clearly the ability of tumor cells to transition to a mesenchymal state contributes to cell invasion, drug resistance, and cell survival in response to the stresses of the primary tumor environment and upon systemic dissemination ^4, 5, 6^. Following dissemination, return to an epithelial state is consistent with an enhanced ability of cells to overcome dormancy and undergo metastatic outgrowth ^3, 7^. In addition to these tumor cell autonomous effects of epithelial-mesenchymal plasticity (EMP) that take place at various steps in the metastatic process, differential EMP conversion rates within the primary tumor contribute to dynamic paracrine relationships between tumor cell populations of varying epithelial or mesenchymal status. We recently termed this concept, epithelial-mesenchymal heterogeneity (EMH) ^8, 9^. An understudied concept of EMP and EMH includes characterization of the epithelial phenotype that remerges after carcinoma cells transition to and from a mesenchymal state ^10^. Herein we sought to address the hypothesis that, following induction of EMT, tumor cells will return to an epithelial state that is similar but critically unique from their original epithelial phenotype.

A key aspect of secondary tumor formation is the ability of disseminated cells to alter their surrounding extracellular matrix to create a niche that is capable of supporting tumor initiation within the context of a normal organ ^11^. Tissue transglutaminase 2 (TGM2) is a crosslinking enzyme, which similar to other transglutaminases catalyzes protein crosslinking via formation of isopeptide bonds between the epsilon-amino group of a lysine and the gamma-carboxamide group of a glutamine ^12^. The ability of TGM2 to crosslink various extracellular matrix proteins including laminin, collagen and fibronectin (FN) is strongly linked to fibrosis and cancer ^13, 14^. In addition to functioning as a freely secreted molecules TGM2 and FN have been detected on the surface of extracellular vesicles (EVs) ^15, 16^. The presence of matrix proteins on the surface of EV contributes to the organotrophic delivery of their various molecular cargoes such as the protein and RNA belonging to their cells of origin ^16, 17^. Therefore, the differential makeup of EVs shed by cancer cells contributes to their ability to reprogram distant stromal cells and directly create a metastatic niche.

Herein we demonstrate that TGM2 and crosslinked FN are upregulated on EVs isolated from metastatic breast cancer cells that have undergone EMP. Furthermore, we utilize *in vivo* approaches and a novel 3D culture model of the metastatic niche to establish that the ability of EVs to reprogram pulmonary fibroblasts to induce the growth of breast cancer cells is strongly dependent on the presence and function of TGM2. These results provide rationale for development of TGM2-targeted biomarkers and therapeutics for the diagnosis and treatment of metastatic breast cancer.

## Results

### Global characterization of gene expression following epithelial-mesenchymal plasticity

Our recent studies demonstrate that induction of EMP via a 4-week treatment with TGF-β1 followed a 2 week withdrawal is sufficient to induce metastasis of HER2 transformed mammary epithelial cells (HME2) upon mammary fat pad engraftment ^3^. Subculture of these bone metastases (HME2-BM) resulted in an epithelial cell population that is morphologically indistinguishable from the parental HME2 cells ^3^. To characterize these two epithelial populations, we performed RNA sequencing analyses on the parental HME2 cells, the purely mesenchymal population that resulted immediately following TGF-1 treatment β (HME2-TGF-β), and the HME2-BM cells (GSE#115255). Analysis of the gene expression data clearly β indicated that long term TGF-β1 treatment induced a gene expression profile that is characteristic of EMT and very unique from the related epithelial states of the HME2 parental and HME2-BM populations (Fig 1a). However, further analysis revealed a set of genes whose expression does not change during the onset of the mesenchymal phenotype induced by TGF-β1, but only becomes significantly upregulated following metastatic reversion to the secondary epithelial state of the HME2-BM cells (Cluster #1; Fig 1A and 1B). Expression of several genes known to be involved in cancer progression were identified by this method, including HMGA2, FHL1, SLC2A3, ADAMTS1, UPP1 and TGM2. Use of qRT-PCR and immunoblot analyses to confirm our RNA sequencing analyses demonstrated robust increases in transglutaminase-2 (TGM2) mRNA and protein levels in HME2-BM cells as compared to the HME2-parental and TGF-β1 treated populations (Fig 2A-2B). In contrast to this mode of TGM2 regulation following completion of EMP, the traditional EMT-associated genes, E-cadherin and fibronectin, showed patterns of robust down and upregulation during TGF-β1-induced EMT, respectively (Fig 2B). Following EMP, expression of E-cadherin returned to levels that were indistinguishable from the parental HME2 cells, but FN levels remained elevated (Fig 2B). Since TGM2 is known to crosslink FN in the extracellular matrix, we analyzed breast cancer patient survival times based expression of FN and TGM2^18^. Similar to what we have previously demonstrated for FN expression alone, patients expressing high levels of both FN and TGM2 demonstrate decreased survival compared to those patients expressing low levels of these two genes (Fig 2C; ^9, 19^). These data suggest that TGM2 in conjunction with FN are clinically relevant markers of EMP whose enhanced expression within the primary tumor is consistent with metastatic disease progression.

**Figure 1.**
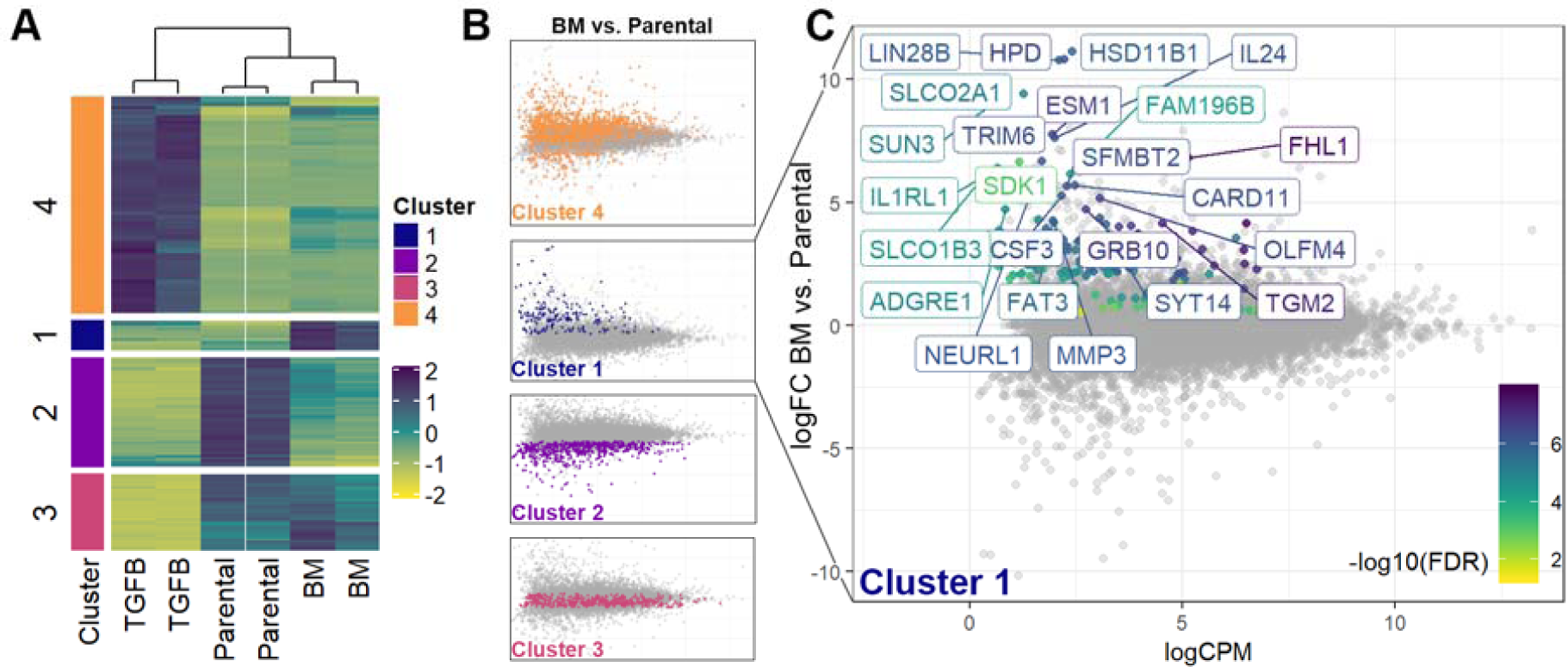
Global characterization of gene expression following metastatic EMP. (A) Dendrogram showing analysis of duplicate RNA sequencing analyses conducted on HME2 cells left untreated (Parental), treated with TGF-β1 for 4 weeks (TGFB) to induce a mesenchymal state, and TGF-β1 treated HME2 cells subcultured from a bone metastasis that formed subsequent to mammary fat pad engraftment (BM). As described in the materials and methods gene expression changes were divided into 4 clusters based on differential expression between the three groups. (B) Cluster 1 was defined as genes whose expression did not change during TGF-β1-induced EMT but were significantly upregulated in the HME2-BM cells as compared to the HME2-parental cells. (C) Identification of an EMP signature of genes whose expression was significantly increased only after induction and metastatic reversion of EMT.

**Figure 2.**
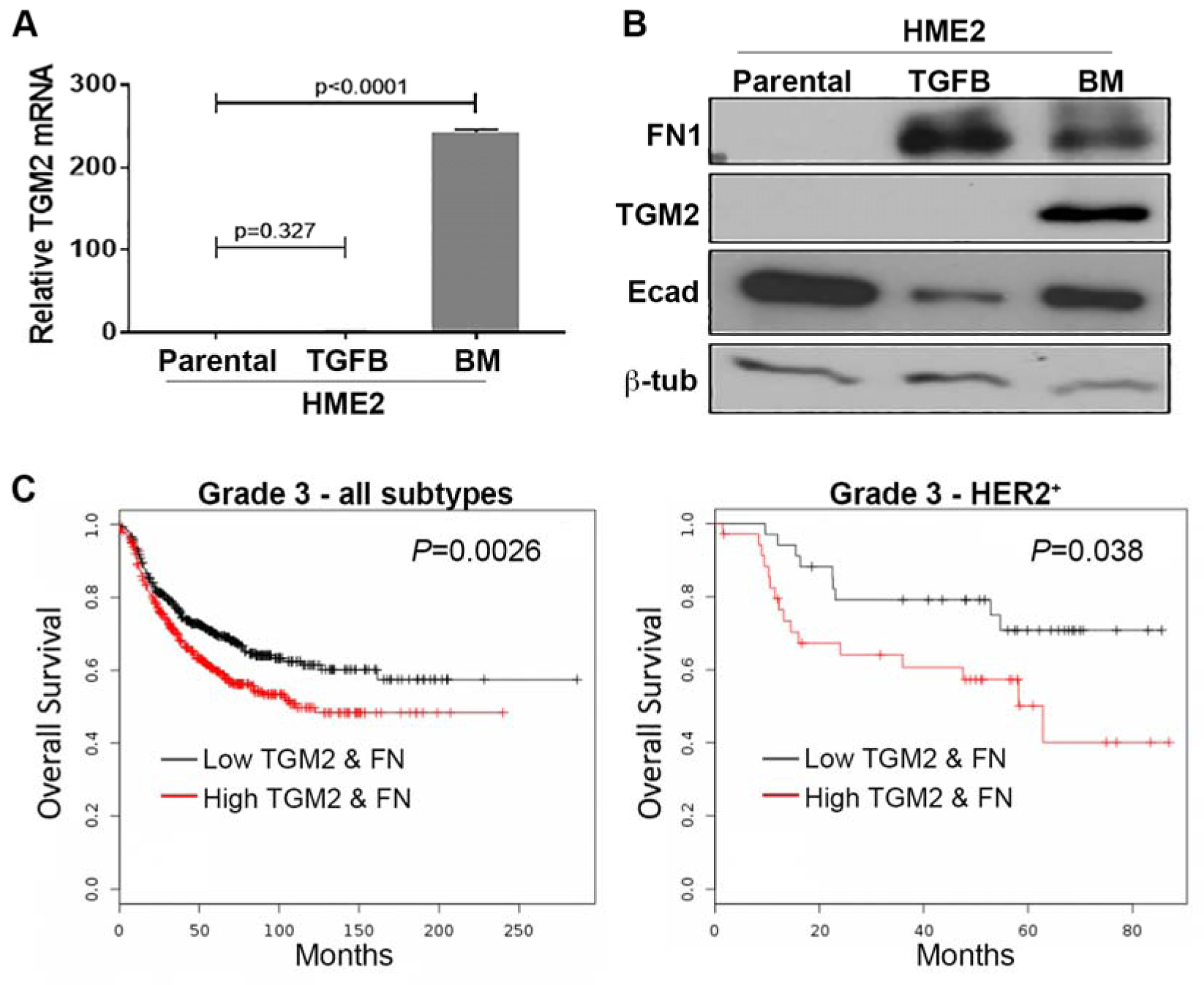
Transglutaminase 2 expression is associated with decreased patient survival. (A) Transcript levels for *TGM2* in HME2 Parental, TGF-β1 treated (TGFB) and bone metastases (BM) were quantified using qRT-PCR. Data are expressed relative to HME2-parental cells and are the mean ±SE of three independent experiments resulting in the indicated p values. (B) Immunoblot analyses for TGM2, FN1, and E-cadherin (Ecad) in HME2 Parental, TGF-β1 treated (TGFB) and bone metastases (BM). Expression of β-tubulin served as a loading control. Data are representative of at least three independent experiments. (C) Comparison of overall survival between patients bearing grade 3 tumors expressing levels of TGM2 and FN above (high) or the below (low) the mean. Survival curves were analyzed via a logrank test resulting in the indicated P values. .

### Transglutaminase-2 promotes breast cancer metastasis

To determine if TGM2 is functionally involved in metastasis, we depleted its expression in the HME2-BM cells and engrafted these cells onto the mammary fat pad of NRG mice (Fig 3A-3B). Depletion of TGM2 had a minimal effect on primary tumor growth but inhibited pulmonary metastasis and promoted overall and metastasis-free survival (Fig 3C-G, Supplementary Fig 1A). To examine the sufficiency of TGM2 in promoting disease progression we overexpressed it in the parental HME2 cells and similarly assessed *in vivo* tumor growth and metastasis (Fig 3H). In contrast to depletion of TGM2 in the HME2-BM cells, overexpression of TGM2 in HME2 cells did significantly increase the growth rate of primary tumors (Fig 3I, Supplementary Fig 1B-C). Moreover, we did observe pulmonary metastasis in TGM2 overexpressing HME2 cells, a result we have yet to observe from parental HME2 tumors in this and other studies (Fig 3I-L, Supplementary Fig 1B-D; ^3, 20^).

**Figure 3.**
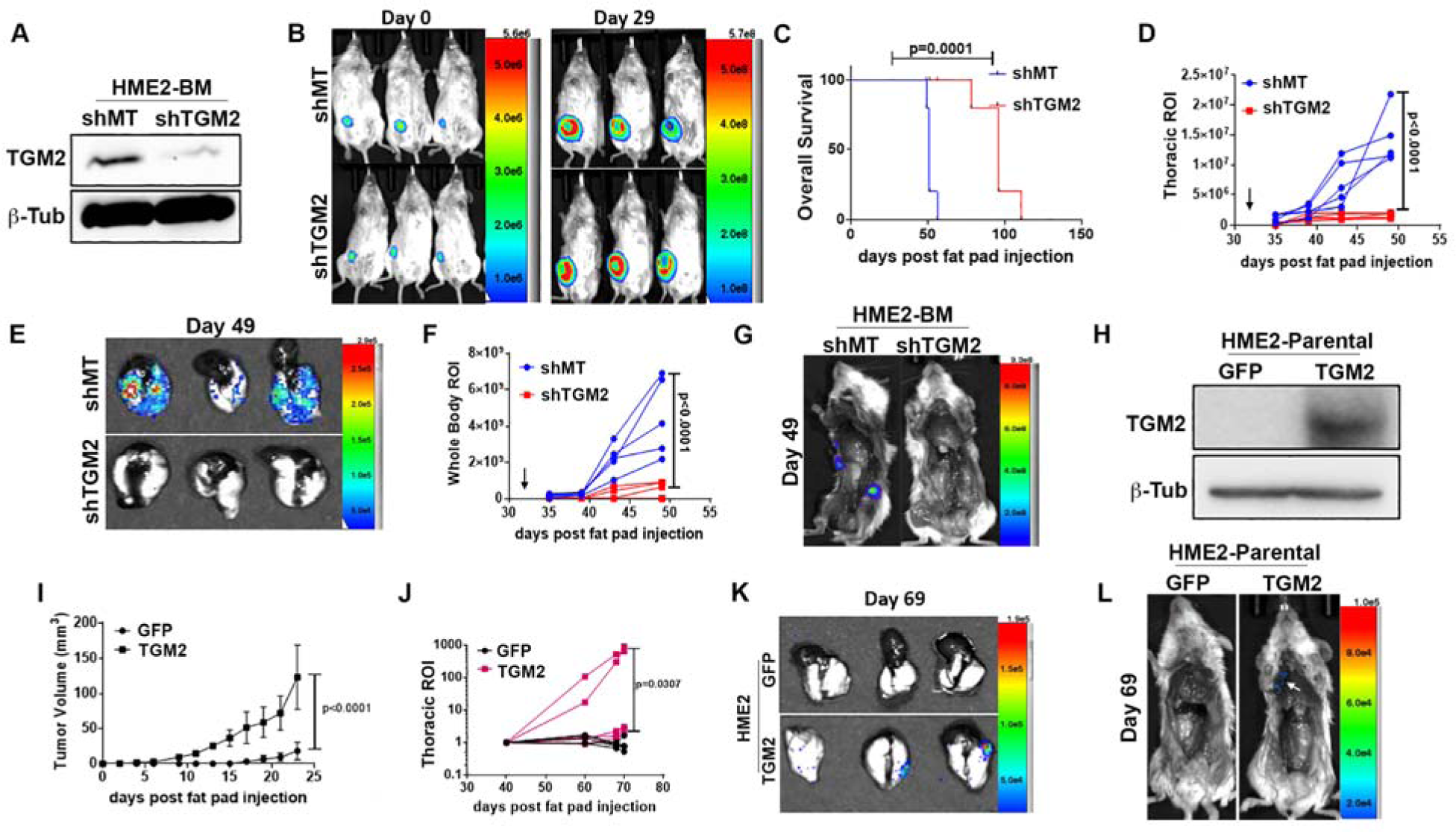
TGM2 drives metastasis. (A) Immunoblot analyses for TGM2 in HME2-BM cells expressing TGM2-targeted shRNAs (shTGM2) or an empty vector (shMT) as a control. Expression of β-tubline (β-Tub) served as a loading control. (B) Cells described in panel A were engrafted onto the mammary fat pad of two separate groups of mice. Bioluminescent images were taken immediately after engraftment (Day 0) and 29 days later (Day 29). (C) Comparison of overall survival between control (shMT) and TGM2 depleted (shTGM2) HME-BM tumor-bearing mice. (D-G) Primary mammary tumors were removed 32 days after engraftment (arrows in D and F) and mice were sacrificed on day 49. Bioluminescent intensity measurements of thoracic regions of interest (ROI; panel D) and whole-body ROI (panel F) of control (shMT) and TGM2-depleted (shTGM2) HME2-BM tumor-bearing mice. Upon necropsy lungs were removed and imaged (panel E) separate of the body (panel G) to visualize pulmonary and extrapulmonary metastases. (H) Immunoblot analyses of TGM2 overexpression in HME2 cells. Expression of GFP was used as a control and β-tubulin (β-Tub) was assessed as a loading control. (I) Primary tumor growth of control (GFP) and TGM2-overexpressing (TGM2) HME2 tumor-bearing mice was quantified by caliper measurements. (J) Bioluminescent intensity measurements of thoracic regions of interest (ROI) of control (GFP) and TGM2-overexpressing (TGM2) HME2 tumor-bearing mice. (K and L) Upon necropsy, the lungs (panel K) and whole mouse (panel L) were imaged separately to visualize pulmonary and extrapulmonary metastases. Data in panels C,D,F,I and J are the mean ±SE or individual values of 5 mice per group resulting in the indicated P values. Data in panel J are the natural log of the bioluminescent radiance for each mouse.

To validate these observations in an additional model of breast cancer metastasis, we deleted TGM2 using a CRISPR-mediated gene editing approach in the highly metastatic 4T1 cells (Fig 4A). We have previously established that the 4T1 undergo robust EMP during tumor growth and metastasis ^3, 21^. Consistent with these studies and our previous results herein, the 4T1 cells expressed readily detectable levels of TGM2 and its deletion hindered 4T1 outgrowth under single cell 3D culture conditions (Fig 4B and Supplementary Fig 2A). Deletion of TGM2 also inhibited primary tumor growth and the pulmonary and extrapulmonary metastasis of the 4T1 cells (Figure 4C-F and Supplementary Fig 2B). These results are consistent with the notion that TGM2 is both necessary and sufficient to promote breast cancer metastasis ^22^.

**Figure 4.**
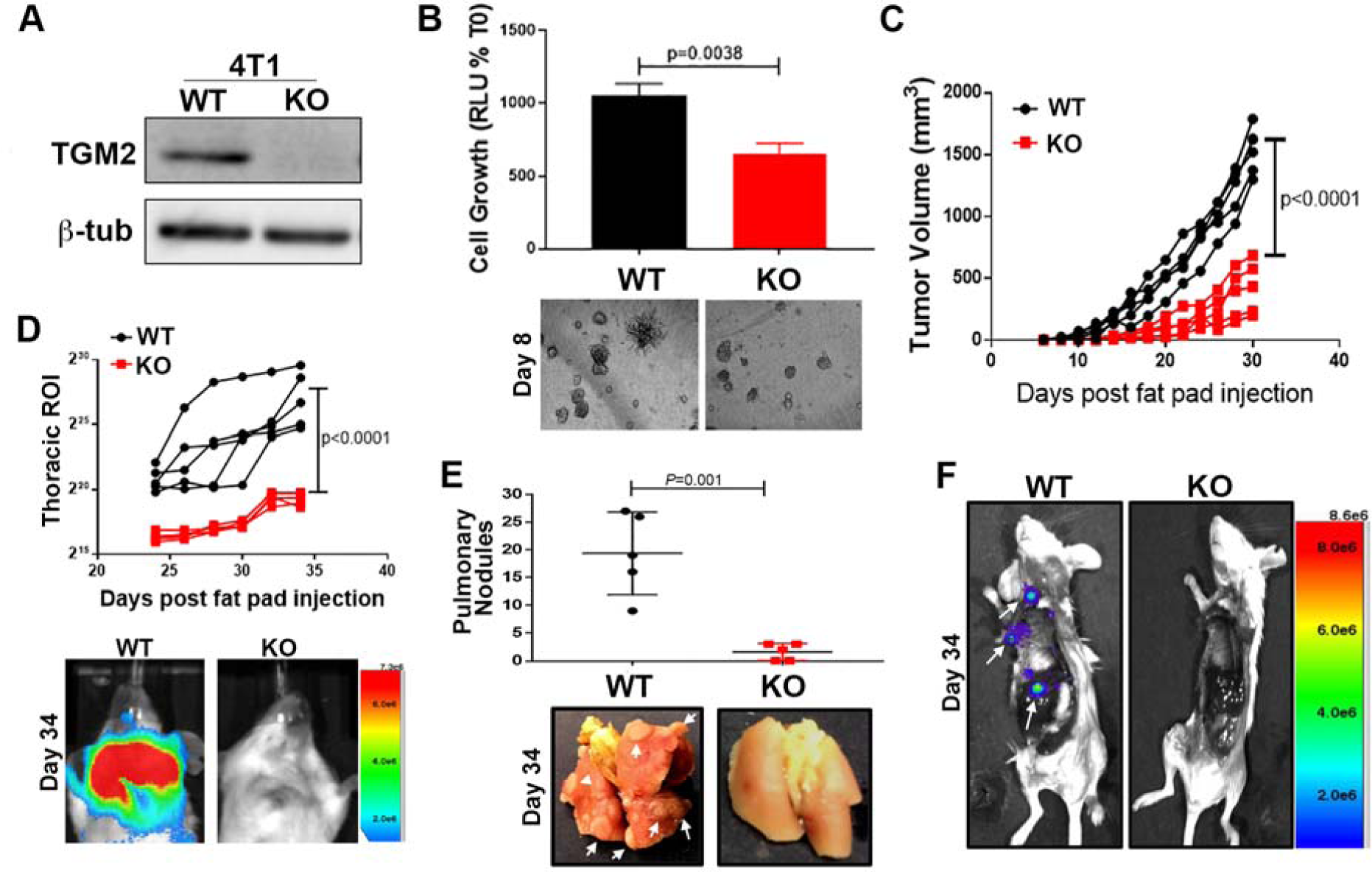
Deletion of TGM2 inhibits metastasis. (A) Immunoblot for TGM2 in control, wildtype (WT) and *Tgm2* deleted (KO) 4T1. Expression of β-tubulin (β-tub) was used as a loading control. (B) Control (WT) and *Tgm2* deleted (KO) 4T1 cells were seeded under single cell 3D culture conditions. Initiation of 3D outgrowth was quantified by bioluminescence. Data are normalized to the plated values and are the mean ±SD of three independent analyses resulting in the indicated P-value. (below) Representative brightfield images of each 3D culture. (C) Control (WT) and *Tgm2* deleted (KO) 4T1 cells were engrafted onto the mammary fat pad and primary tumor growth was quantified by caliper measurements. Data are of individual mice taken at the indicated time points, resulting in the indicated *P*-value. (D) Quantification of bioluminescent radiance from the pulmonary regions of interest (ROI) at the indicated time points. (below) Representative thoracic bioluminescent images of control 4T1 (WT) *Tgm2* deleted (KO) tumor-bearing mice. (E) Upon necropsy, the numbers of metastatic pulmonary nodules were quantified from control (WT) and *Tgm2* deleted (KO) 4T1 tumor-bearing mice.(below) Representative gross anatomical images of lungs from these groups. (F) Upon necropsy the lungs and primary tumors of mice bearing control (WT) and *Tgm2* deleted (KO) 4T1 tumors were removed and the carcasses were immediately to visualize extra pulmonary metastases (arrows). A representative mouse from each group is shown. For panels D and E data are the mean ±SE of 5 mice resulting in the indicated P values.

### Transglutaminase-2 promotes fibronectin crosslinking on the surface of extracellular vesicles

Previous studies indicate that TGM2 and FN are present in extracellular vesicles derived from cancer cells ^15, 23^. We therefore isolated EVs using a 200 nM filter cutoff from non-metastatic and metastatic breast cancer cells and conducted nanoparticle tracking analysis and transmission electron microscopy to validate vesicle isolation (Fig 5A and Supplementary Fig 3A). These EV fractions were further analyzed by immunoblot for expression of TGM2 and the crosslinked status of FN (Fig 5B). Crosslinked FN dimers were only observed in vesicles derived from the metastatic HME2-BM cells that had undergone EMP, not in vesicles from the non-metastatic HME2 parental cells (Fig 5B). Genetic depletion of TGM2 or pharmacological inhibition of its activity using the small molecule NC9, inhibited FN crosslinking on EVs (Fig 5B, Supplementary Fig 3B). Finally, overexpression of TGM2 was sufficient to induce FN dimerization on EVs and this could readily inhibited by NC9 (Fig 5B, Supplementary Fig 3B). To further investigate the hypothesize that crosslinked FN progresses to a fibrillar form on the surface of EVs we conducted confocal microscopy on nonpermeabilized EVs using a fibular FN specific antibody (Fig 5C). Results from this approach are consistent with the conclusion that fibrillar FN exists on the surface on EVs in a TGM2 dependent manner (Figure 5B-D). Taken together, these data indicate that TGM2-mediated crosslinking promotes FN fibrillogenesis on the surface of EVs derived from metastatic breast cancer cells that have undergone EMP.

**Figure 5.**
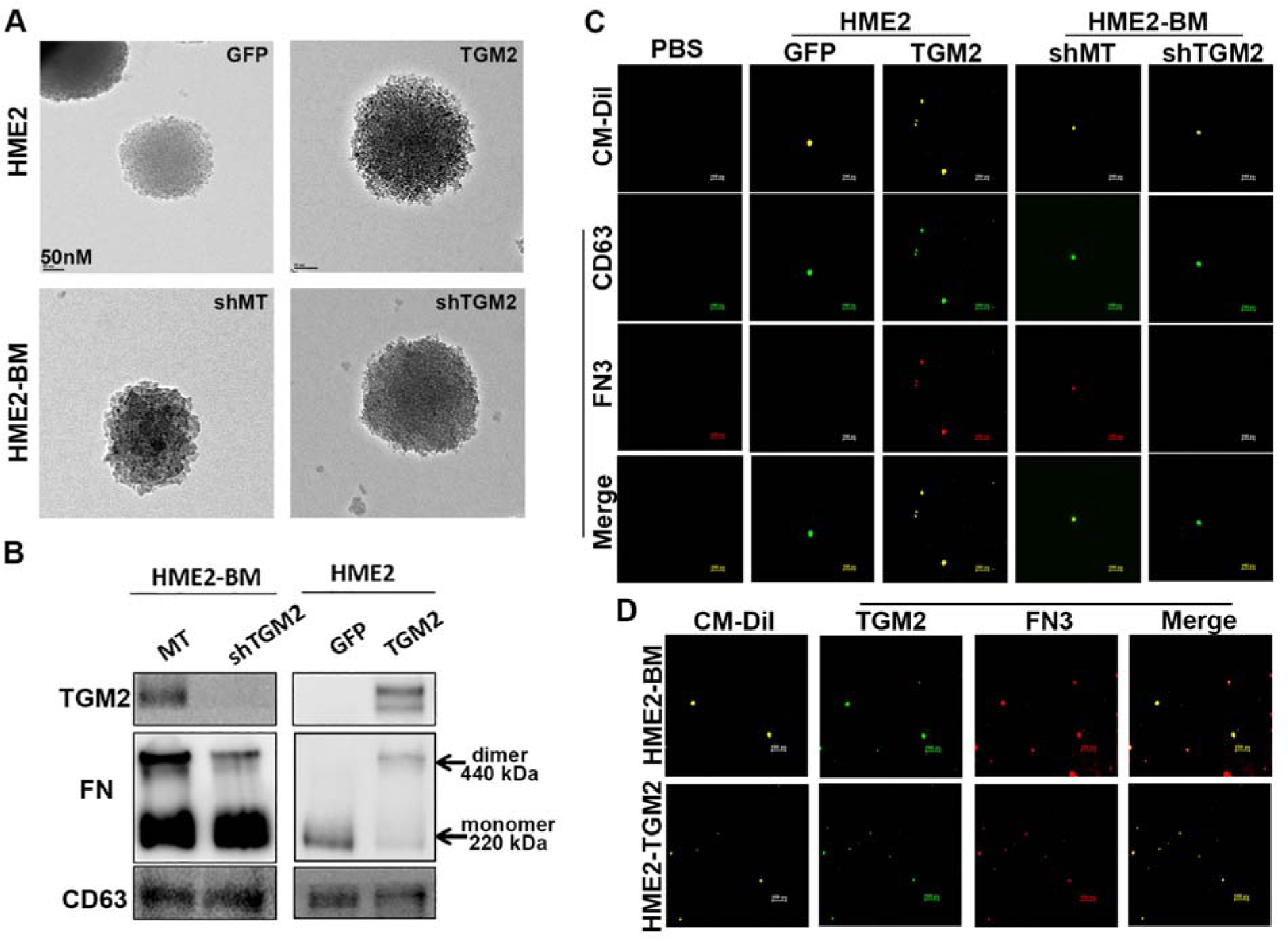
Transglutaminase-2 promotes fibronectin crosslinking on the surface of extracellular vesicles. (A) Transmission electron micrographs of extracellular vesicles derived from control (GFP) and TGM2 overexpressing HME2, and control (shMT) and TGM2 depleted (shTGM2) HME2-BM cells. (B) Immunoblot analysis of EVs derived from the HME2 and HME2-BM cells described in panel A. Differential expression of TGM2 was verified in these EV lysates and correlated with covalent linkage of FN dimers that are insensitive to reducing conditions of the SDS-PAGE. CD63 served as a loading control. (C) Extracellular vesicle preparations derived from the cell types described in panel A were stained with CM-Dil (yellow) to verify the presence of a lipid containing particles. These preparations were also stained with antibodies specific for CD63 (green) and FN3 (red) and imaged using confocal microscope. The green (CD63) and red (FN3) channels were merged. A blank control sample (PBS) stained with the CM-Dil and appropriate secondary antibodies is also shown. (D) Extracellular vesicles derived from HME2-BM and HME2-TGM2 cells were stained with CM-Dil (yellow), and antibodies specific for TGM2 (green) and FN3 (red) and imaged using a confocal microscope. The green (TGM2) and red (FN3) channels were merged. Scale bars on panels C and D are 500 nm.

### Tensin-1 is required for fibronectin fibrillogenesis on extracellular vesicles

Our recent phospho-mass spectrometry analyses indicate increased phosphorylation of Tensin-1 (TNS1) in EVs derived from the serum of breast cancer patients as compared to healthy individuals ^24^. Tensin-1 plays an important role in FN fibrillogenesis by binding to and promoting clustering of ECM-bound integrins ^25^. Consistent with these patient data, TNS1 was readily detectable in EV fractions, and was markedly dependent on the expression of TGM2 (Fig 6A, Supplementary Fig 4). In contrast, depletion of Tns1 in the 4T1 cells did not affect TGM2 presence on EVs or FN dimerization (Fig 6B and 6C). Despite the fact that Tns1 was difficult to detect in the 4T1 model and its depletion did not affect TGM2, use of the fibrillar FN specific antibody in conjunction with immunoelectron microscopy indicated that FN fibrillogenesis was not achieved on EVs in the absence of Tns1 (Fig 6D). Functionally, depletion of Tns1 inhibited the outgrowth of the 4T1 cells under single cell 3D culture conditions (Fig 6E). Upon orthotopic engraftment, Tns1 depletion had no effect on primary tumor growth but did significantly inhibit pulmonary metastasis (Fig 6F-6H). Together with our previous work, these data suggest that TGM2 mediates the presence of Tns1 in tumor-derived EVs, leading to FN fibrillogenesis and promotion of metastasis.

**Figure 6.**
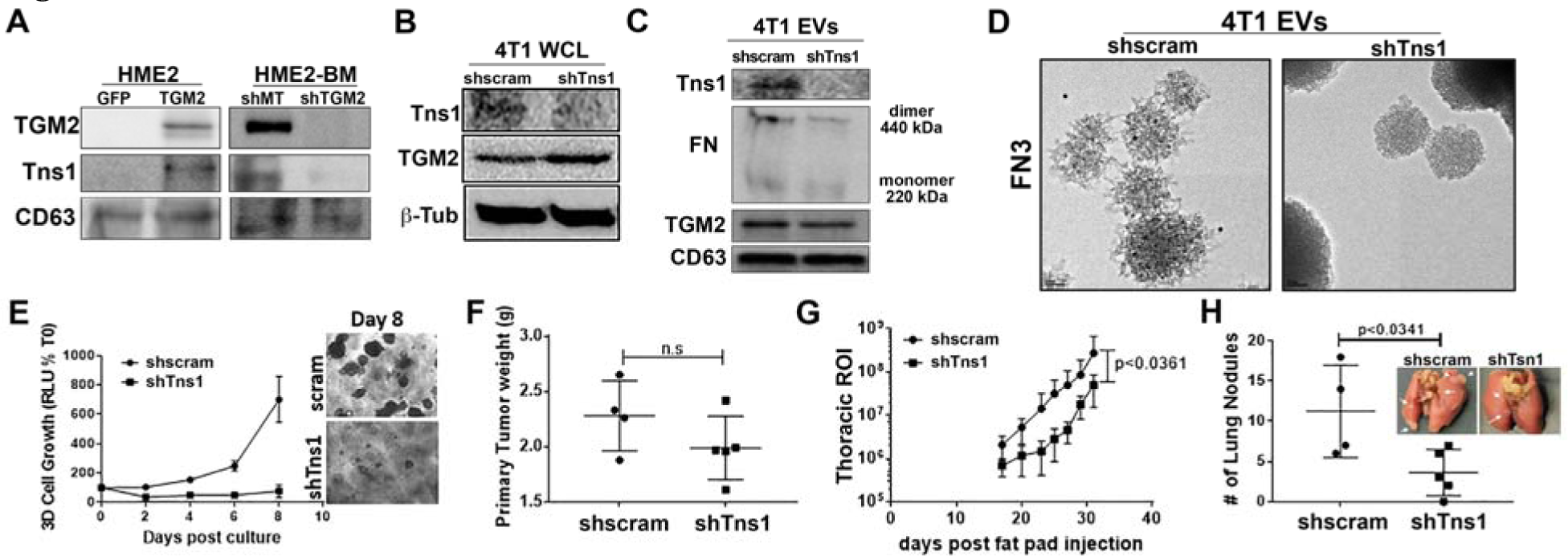
TNS1 is required for fibronectin fibrillogenesis on EVs. (A) Immunoblot analyses of EVs collected from control (GFP) and TGM2 overexpressing HME2, and control (shMT) and TGM2 depleted (shTGM2) HME2-BM cells. Lysates from these EVs were assessed for levels of TGM2, Tns1, levels of CD63 served as a loading control. (B) Immunoblot analysis of whole cell lysates (WCL) derived from control (shscram) and Tns1 depleted (shTns1) 4T1 cells were analyzed for expression of Tns1, TGM2. β-Tubulin (β-Tub) served as a loading control. (C) Immunoblot analyses of EVs derived from control (shscram) and Tns1 depleted (shTns1) 4T1 cells. Lysates from these EVs were assessed for levels of TGM2, Tns1, levels of CD63 served as a loading control. (D) The fibillar FN specific antibody (FN3) antibody was used in conjunction with immuno-electron microscopy to analyze nonpermeabilized EVs derived from control (shscram) and Tns1 depleted (shTns1) 4T1 cells. (E) Control (shscram) and Tns1 depleted (shTns1) 4T1 cells were grown under single cell 3D culture conditions. Cellular outgrowth was quantified by bioluminescence at Day 8. Data are normalized to the plated values and are the mean ±SD of three independent analyses resulting in the indicated P-value. (F) Control (shscram) and Tns1 depleted (shTns1) 4T1 cells were engrafted onto the mammary fat pad and the resultant primary tumors were removed and weighed upon necropsy. (H) Quantification of bioluminescent radiance from the thoracic regions of interest (ROI) at the indicated time points. (H) Upon necropsy, the numbers of pulmonary metastatic nodules were quantified. (Inset) Corresponding gross anatomical views of lungs from control (shscram) and Tns1 depleted (shTns1) 4T1 tumor-bearing mice. Data F-H are mean ±SE values from five mice per group resulting in the no significance (n.s.) or the indicated P-values.

### Transglutaminase-2 promotes extracellular vesicle-mediated metastatic niche formation

To focus on the specific role of EVs in TGM2 and TNS1-mediated metastasis, we developed a coculture model of the pulmonary niche. To do this, human pulmonary fibroblasts (HPF) were grown to confluence on our recently described tessellated polymeric scaffolds to create a three-dimensional platform for subsequent coculture with tumor cells (Fig 7A; ^9^). The HPFs were treated with EVs derived from control, TGM2-deleted, or TNS1-depleted 4T1 cells for three weeks in attempts to modulate the growth environment (Fig 7A). Following pretreatment with these EVs, responder cells (MCF10-Ca1a cells expressing a firefly luciferase-dTomato fusion protein) were seeded onto the HPFs (Fig 7A). Using this approach, we observed that HPFs pretreated with WT 4T1-derived EVs significantly enhanced the growth of responder cells as compared to untreated HPFs (Fig 7B and 7C). Moreover, depletion of TGM2 or TNS1 significantly reduced the ability of EVs to induce the growth supportive phenotype of the HPFs (Fig 7B and 7C).

**Figure 7.**
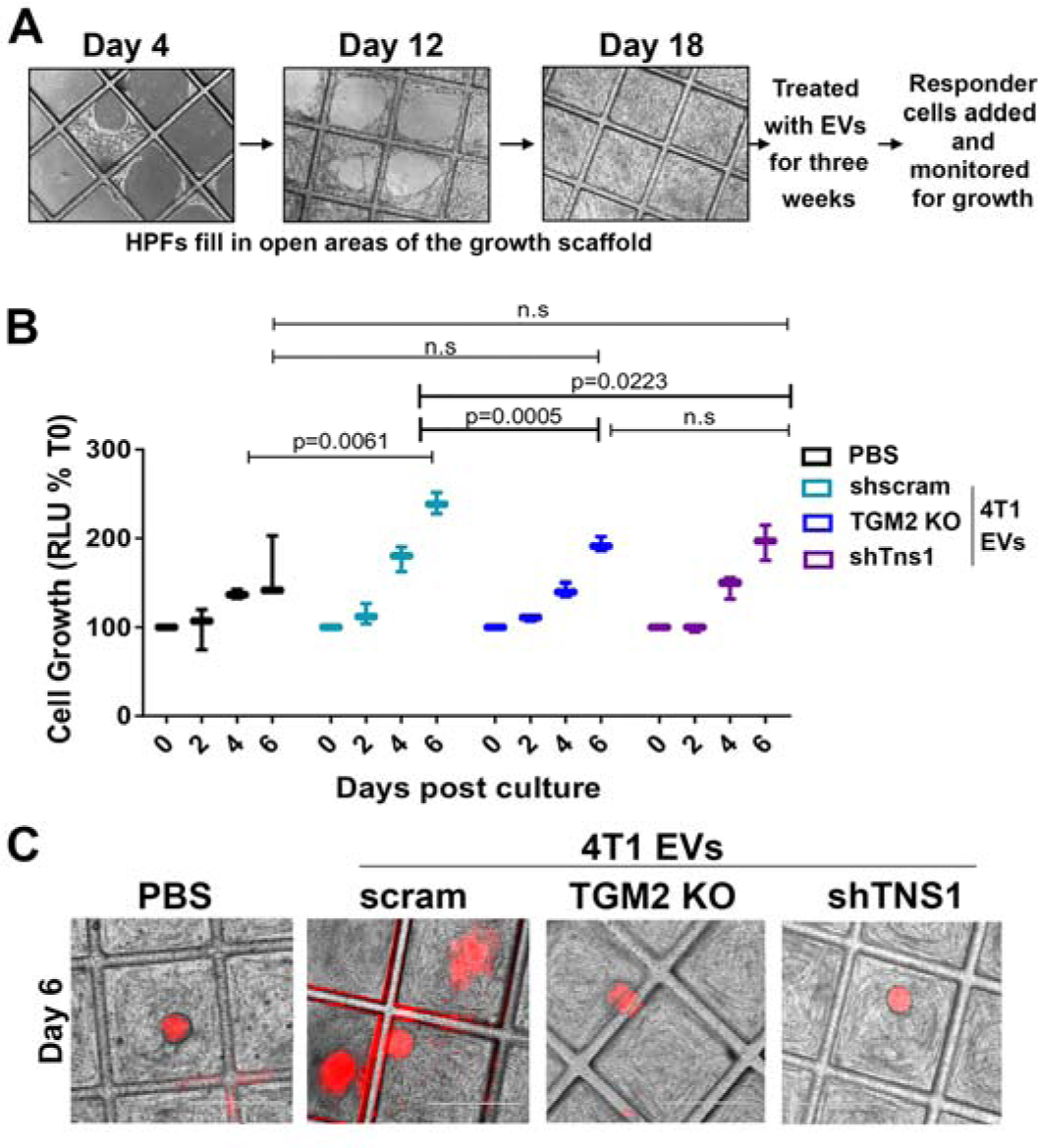
TGM2 promotes EV-mediated pulmonary niche formation. (A) Schematic illustration of the 3D niche assay. Human pulmonary fibroblasts were cultured on 3D scaffolds as described in methods and allowed to fill the open space of tessellated polymeric scaffolds. These cultures were treated with exosomes for another 3 weeks at which point MCF10-Ca1a responder cells stably expressing a firefly luciferase-dTomato fusion protein were added to the culture. (B) Growth of labeled Ca1a cells was longitudinally quantified by bioluminescence at the indicated time points. Data are mean relative luminescence units (RLU), normalized to the time zero (T0) reading, ±SE for three independent experiments completed in triplicate resulting in the indicated P values. (C) Representative fluorescent and brightfield merged images showing Cala cell colonies (red) growing on the HPFs.

Next, we again utilized our 3D coculture system to examine the ability of EVs from HME2 cells to promote the growth of responder cells. Indeed, HPFs treated with EVs derived from parental HME2 cells failed to enhance the growth of responder cells, but this could be drastically enhanced by overexpression of TGM2 (Fig 8A). Conversely, HME2-BM-derived EVs promoted responder cell growth, and this was prevented upon depletion of TGM2 in the EVs (Fig 8B). Importantly, prevention of FN crosslinking via treatment of EV-producing cells with NC9 also prevented the ability of the resultant EVs to induce the growth supportive phenotype of the HPFs (Fig 8A-B). To confirm that EVs derived from metastatic breast cancer cells promote metastatic niche formation, NSG mice were treated with WT and TGM2-depleted EVs via intraperitoneal injections for 3 weeks. These mice were then subsequently given tail vein injections of the bioluminescent MCF10-Ca1a responder cells. Consistent with our 3D coculture results, *in vivo* administration of HME2-BM-derived EVs enhanced the pulmonary colonization of responder cells (Fig 8C-8E). Importantly, this effect was abolished when TGM2 was depleted from the administered EVs (Fig 8C-8E). Altogether, these results suggest that EVs derived from metastatic breast cancer cells utilize the aberrant presence of TGM2 to educate pulmonary fibroblasts cells to form a pulmonary niche more suitable for metastatic colonization.

**Figure 8.**
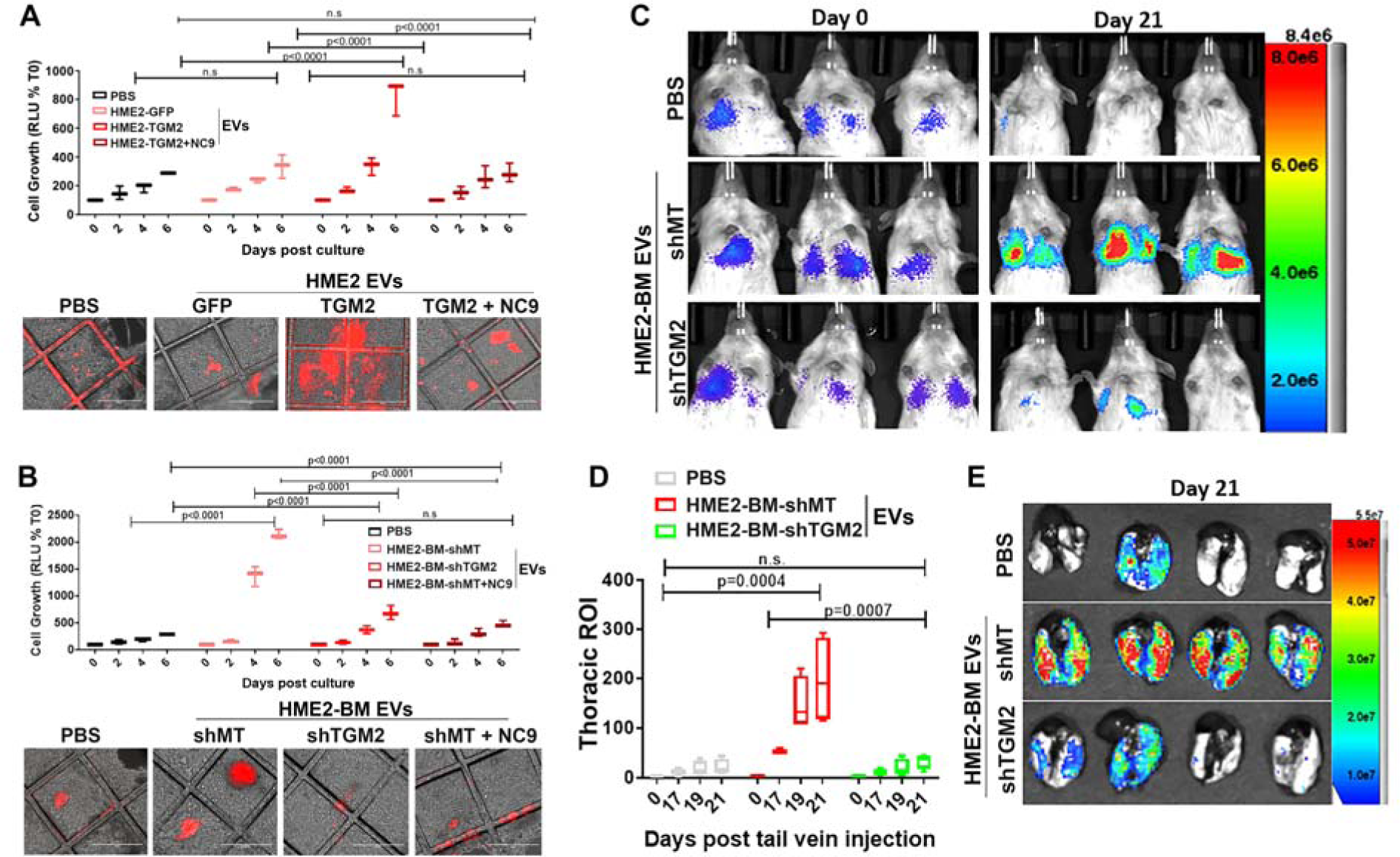
TGM2 enhances the ability of EVs to increase pulmonary colonization. (A-B) Three dimensional cultures of human pulmonary fibroblast were treated with the indicated EVs as described in the materials and methods. MCF10-Ca1a cells stably expressing a firefly luciferase-dtomato fusion protein were subsequently added to these cultures and their growth longitudinally quantified by bioluminescence at the indicated time points. Data are mean relative luminescence units (RLU), normalized to the time zero (T0) reading, ±SE for three independent experiments completed in triplicate resulting in the indicated P values. Representative fluorescent and brightfield merged images showing Cala cell colonies (red) growing on the HPFs. (C) Mice were pretreated with indicated EVs via intraperitoneal injections for 3 weeks prior to tail vein administration of the labeled Ca1a described in panel A. Bioluminescent images of representative mice taken immediately (Day 0) and 21 days (Day 21) following tail vein injection of the Ca1a cells. (D) The mean (±SE) bioluminescence values of thoracic regions of interest (ROI) taken at the indicated time points. Data are normalized to the injected values, n=4 mice in each group resulting in the indicated P value. (E) Mice were sacrificed at day 21 and upon necropsy lungs were imaged *ex-vivo* using bioluminescence to visualize pulmonary tumor formation.

## Discussion

Several recent studies from our lab and others indicate that dynamic induction and reversion of EMP drives tumor cell heterogeneity and supports more efficient completion of several steps of the metastatic process ^8, 9, 26^ Herein we present a global characterization of gene expression changes that only .emerge after cells complete EMP. This approach led to the discovery of TGM2 upregulation as a marker of EMP. We went on to validate the necessity and sufficiency of TGM2 to promoting metastasis and explored the impact of TGM2 on FN fibrilization and function of EVs in producing a metastatic niche.

TGM2 has previously been linked to EMT and, in contrast to our observations, its expression has been observed in cells with a mesenchymal morphology ^27^. These data clearly suggest model dependent changes in gene expression that occur during EMP. For instance, TGF-β stimulation of the HME2 model results in a mesenchymal phenotype which, upon cessation of cytokine treatment, is fully capable of transitioning back to an epithelial morphology indistinguishable from the original cells ^28^. In contrast, different cell models or differences in EMT-inducing stimuli may result in EMT events that have a spectrum of gene expression and morphological reversion capabilities. Indeed, using the HME2 model we recently established that chronic inhibition of HER2 kinase activity with the drug Lapatinib results in an EMT that does not revert upon removal of the drug ^28^. Therefore, some EMT events may revert to an extent that includes upregulation of TGM2, or other genes, but may not include an overt morphologic change back to an epithelial state. This concept would serve to explain why certain cell models that have a stable mesenchymal morphology are fully capable of completing metastasis, while others fail to complete the latter stages of metastasis. Overall, our findings in the HME2 model highlight the importance of defining EMT and MET by quantitative changes in gene expression or other metrics, not by morphologies that manifest under two-dimensional culture conditions.

Additional explanation in regard to the role of TGM2 in EMT may also stem from its diversity in function both inside and outside the cell ^29^. Here, we focused on characterizing the systemic influence of TGM2 containing EVs in the development of niches with increased capacity to support tumor cell seeding ^30^. Both FN and TGM2 have been observed in EVs derived from breast cancer cells ^15, 31^. In this study, we demonstrate the necessity and sufficiency of TGM2 to generate fibrillar FN on the surface of EVs, an event that fosters pulmonary niche formation and supports the colonization of breast cancer cells. These functional data are supported by our patient analyses that suggest TGM2 and FN function together to drive tumor progression and our previous studies noting increased phosphorylation of TNS1 in EVs derived from breast cancer patients ^24^.

Extracellular vesicles derived from cancer cells contain multiple biologically active molecules including proteins, nucleic acids, and lipids that can alter local stroma to create a disease supportive microenvironment. Our studies utilize a novel, tessellated 3D coculture platform to establish that TGM2 is required for EV-mediated modification of pulmonary fibroblasts. However, important questions remain in regard the precise mechanism by which TGM2 and fibular FN participate in this process. Comprehensive analyses are required to characterize changes in the EV components upon TGM2 depletion and/or inhibition in cells that have undergo EMP ^15^. Moreover, the presence of fibrillar FN on the surface of EVs could drastically alter the amount and route of EV internalization into pulmonary fibroblasts, leading to pathologic alteration of fibroblast gene expression and function. Finally, delivery of TGM2 and FN into the stromal pulmonary microenvironment maybe sufficient to alter the existing matrix into a more tumor permissive state. Studies in the lab are currently ongoing to better delineate such mechanisms by which the presence of TGM2 alters EV function.

Clinically, our data suggest that attempts to quantify systemic changes in TGM2 activity as a bodily fluid-based biomarker could be augmented by preparation of EV fractions prior to these analyses ^32, 33^. Finally, our studies support the notion that targeted inhibition of TGM2 will limit disease progression by preventing not only local fibrotic reactions but also through inhibition of distant metastatic niche formation. Overall, our data present comprehensive characterization of EMP and illustrate a novel impact of this process on EV function and metastatic progression.

## Materials and Methods

### Reagents

Luciferase expressing 4T1 and HMLE cells were transformed via overexpression of Her2 and their mesenchymal variants were described previously ^20, 34^. Ca1a cells were kindly provided by Dr. Fred Miller (Wayne St. University). These cells were cultured in DMEM containing 10% FBS and 1% pen/strep. Stable expression of the firefly luciferase or the firefly luciferase-dTomato fusion protein was achieved through lentiviral transduction and selection in blasticidin or zeocin ^9^. Manipulation of TGM2 expression was achieved through lentiviral-mediated transduction of TRCN0000000239, TRCN0000000241, or a scrambled control shRNA, empty plko.1 vector (GE Dharmacon, Lafayette, CO). The dimeric CRISPR RNA-guided Fokl nuclease and Csy4-based multiplex gRNA expression system as previously described was used to generate the TGM2 knockout 4T1 cell line ^3^. Full length human TGM2 or GFP as a control were expressed via a lentiviral delivery of pLV (Vector Builder, Santa Clara, CA). Manipulation of Tns1 expression was achieved through lentiviral-mediated transduction of V2LMM_4992 or a scrambled control (vector builder). Stable genomic integration of constructs was selected for using puromycin or hygromycin. Primary human pulmonary fibroblasts were obtained from ATCC and cultured in the recommended fibroblast basal media supplemented with fibroblast growth factor low serum kit (ATCC). The TGM2 small molecule inhibitor (nc-9) was a kind gift from Dr. Jefferey Keillor (University of Ottawa).

### Gene expression analysis

Gene expression data for the HME2 parental, TGF-β1 treated, and HME2-BM cells were extracted from GSE115255 as previously described ^3^. Afterwards, the raw counts were imported to the R environment of statistical computing (v3.6.0). Differentially expressed genes were determined by using edgeR (v3.26.4,^35, 36^) and defined as having a log fold change higher than 2 and adjusted p.value less than 0.05. Then the genes were grouped using unsupervised clustering via k-means, determining the optimal k by the elbow method and plotted using the R packages ComplexHeatmap (v2.0.0,^37^) or ggplot2 (DOI: 10.1007/978-0-387-98141-3). For RT-PCR, total RNA was reverse-transcribed using a cDNA synthesis kit (Thermo Fisher). Semi quantitative real-time PCR was performed using iQ SYBR Green (Thermo Fisher). The following primers were used for the analysis. *TGM2* sense; ATAAGTTAGCGCCGCTCTCC, *TGM2* antisense; CTCTAAGACCAGCTCCTCGG.

### Animal models

All *in vivo* assays were conducted under IACUC approval from Purdue University. Where indicated luciferase expressing HME2 variants (2x10^6^ / 50 µl) were injected into the second mammary fat pad of female 8-week old, NSG mice. Tumors were surgically excised after they achieved size of 900 mm and metastasis was subsequently quantified by bioluminescent imaging using the Advanced Molecular Imager (AMI) (Spectral Instruments, Tucson, AZ). For EV preconditioning experiments, 20ug of total EV protein in 100 µl of sterile PBS was injected intraperitoneally into 3 months old NSG mice every other day for 3 weeks. Mice received 100 µl of sterile PBS in the control group. Luciferase expressing Ca1a (7.5x10^5^ / 100 µl) were injected into the lateral tail vein of EV pretreated female NSG mice. Pulmonary tumor growth was quantified by bioluminescence at the indicated time points.

Luciferase expressing 4T1 cells were resuspended in PBS (50 µl) and orthotopically engrafted onto the second mammary fat pad of 4 weeks old Balb/c mice (2.5x10^4^ cells/mouse) (Jackson Labs, Bar Harbor, ME). Primary tumor growth and metastasis development were assessed via weekly bioluminescent imaging. Upon necropsy, lungs from all animals were removed and fixed in 10% formalin and dehydrated in 70% ethanol for visualization of pulmonary metastatic nodules and histological analyses.

### Isolation of extracellular vesicles

EVs were isolated from 72 hours, serum free conditioned media of 10^7^ cells (equivalent to four 150 mm dishes) as described previously ^38, 39^. Briefly, conditioned media was centrifuged at 300xg for 10 minutes to remove live cells and then 2000xg for 10 minutes to remove cell debris. This supernatant was filtered through a 0.22µM pore size Millipore filter. Filtered media was concentrated to 1 ml using 3KD MWCO Amicon ultra-15 centrifugal filters, followed by ultracentrifugation at 100,000 g for 2h. The pellet was washed with PBS using ultracentrifugation at 100,000 g for an additional 2h. The pelleted EVs were resuspended either in 3D RIPA buffer for immunoblot experiments or in PBS for biological or imaging experiments. Size distribution and concentration of EVs were analyzed via semiautomated nanoparticle tracking using a NanoSight (NS300) apparatus (Malvern Instruments Ltd.). Samples were diluted to provide counts within the linear range of the instrument (3x10^8^ – 1x10^9^ per ml). Three videos of 1-minute duration were documented for each sample, with a frame rate of 30 frames per second. Using NTA software (NTA 2.3; NanoSight Ltd.), particle movement was analyzed as per manufacturer’s protocol. The NTA software was adjusted to first identify and then track each particle on a frame-by-frame basis.

### Immunological assays

Protein expression on the surface of EVs was examined using whole mount immunostaining staining as described previously ^40^. Briefly, thin formvar/carbon film coated 200 mesh copper EM grid were glow discharged for 30 seconds. EVs were fixed in 1 ml of paraformaldehyde (PFA) for 5 mins. Five-7 µl of fixed EVs were loaded onto the grids and incubated for 10 minutes. The grids were rinsed with 100 µl of PBS three times each for 10 min, then treated with 50 µl of glycine to quench free aldehyde groups for 10 min. The grid was then incubated with 100 µl of blocking buffer (PBS containing 1% BSA) for 30 min and finally incubated with 100 µl of primary antibody (anti-FN3 (1:100)) overnight at 4°C. The following day, the grids were washed with 100 µl of washing buffer (PBS containing 0.1% BSA) five times each for 10 min. Following incubation with secondary antibody conjugated to a 10 nm gold particle (ab39619, abcam) diluted at 1:100 in PBS containing 0.1% BSA for 1hour. Grids were washed with washing buffer five times each for 10 min and 50µl of distilled water twice. Grids were air dried for 15 min and observed via TEM at 200kV.

For confocal microscopy imaging, EVs samples were prepared as described previously ^9^. Briefly, 500 µl of PBS solution containing EVs was incubated simultaneously with 2 µl each of antibodies specific for CD63 (ab217345), TGM2 (Invitrogen CUB 7402) and FN3 (Invitrogen 14-9869-82) for 2 hours at room temperature. Following incubation, the solution was purified by ultrafiltration (50 Kda MWCO) at 600 rpm for 20 min. The filtrate was washed with PBS using ultrafiltration and resuspended in PBS. Next, a mixture of 0.5ul of CM-dil, 1 µl of 2 mg/ml of Alexa Fluor labeled 647 Goat anti-mouse IgG, and 1 µl of 2mg/ml of Alexa Fluor 488 labeled Goat anti-rabbit IgG was added to the EV solution, incubated for 1h with vigorous mixing, and then purified again by ultrafiltration. Finally, the precipitate was resuspended in PBS and added to 35 mm^2^ 1.5H glass coverslip bottom confocal dish and adsorbed for 15 minutes. The dishes were coated with 0.1% polyethylenimine for 15 mins prior to addition of prepared sample. Samples were imaged using Nikon confocal microscope.

For immunoblot analyses, cells and EV fractions were lysed using a modified RIPA lysis buffer containing 50 mM Tris, 150mM NaCl, 0.25% Sodium Deoxycholate, 1.0% NP40, 0.1% SDS, protease inhibitor cocktail, 10mM activated sodium ortho-vanadate, 40 mM β-glycerolphosphate and 20mM sodium fluoride. These lysates were separated by reducing SDS PAGE and probed for TGM2, CD63 (Cell signaling technologies, Danvers, MA), fibronectin, epithelial cadherin (Ecad; BD biosciences, San Jose, CA), or β-tubulin (DSHB, Iowa City, IA).

### 3D scaffold assays

Scaffolds were constructed as described previously ^9^. Uncoated 3D scaffolds were placed in ultralow attachment 24-well dishes. Human pulmonary fibroblasts (100,000) were added to the scaffolds. Cells were fed new media every 5 days for 2-3 weeks. Once the scaffolds were entirely covered with HPFs, they were treated with 5ug of EVs every other day for 3 weeks. Ca1a FF-dTomato cells (50000) were added to the EV-pretreated scaffolds and tracked for growth using bioluminescence and fluorescent imaging at indicated time points.

### Statistical analyses

2-way ANOVA or 2-sided T-tests were used where the data met the assumptions of these tests and the variance was similar between the two groups being compared. P values of less than 0.05 were considered significant. No exclusion criteria were utilized in these studies.

## Acknowledgments

This research was supported in part by the American Cancer Society (RSG-CSM130259) to M. Wendt and the National Institutes of Health (R01CA207751, R01CA232589, and R21AA026675) to M. Wendt and (R00CA198929) to L. Solorio, and the Purdue Center for Cancer Research via an NCI center grant (P30CA023168). We kindly acknowledge the expertise of the personnel within the Purdue Center for Cancer Research Biological Evaluation Core. We also acknowledge the use of the facilities within the Bindley Bioscience Center, a core facility of the NIH-funded Indiana Clinical and Translational Sciences Institute.

## Supplementary Figures

**Supplementary figure 1:**
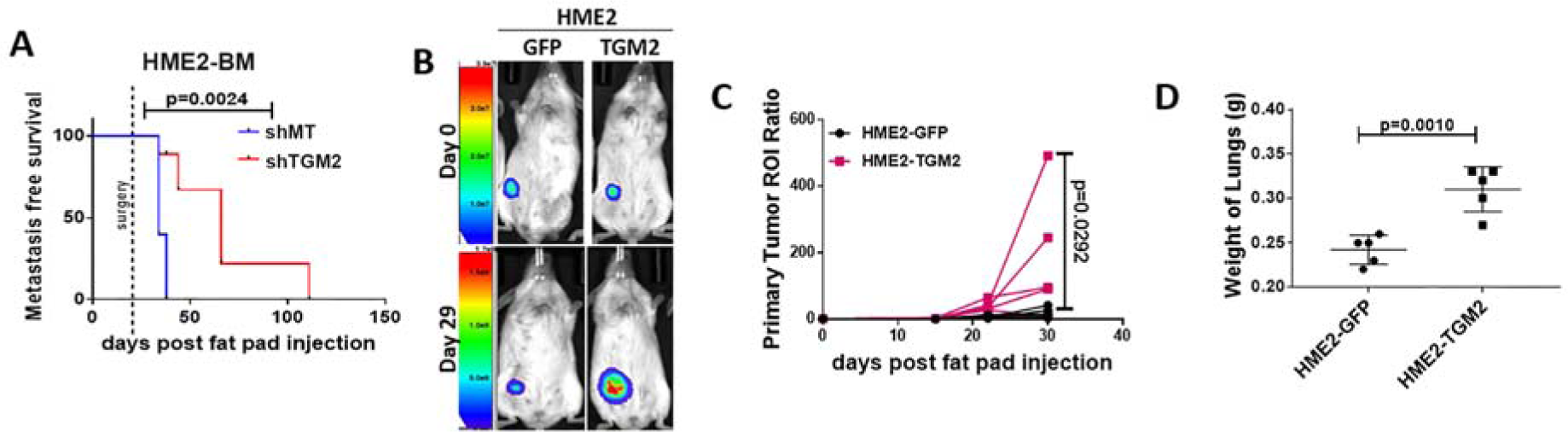
TGM2 expression promotes metastasis and decreases survival. (A) Metastasis free survival analysis of mice orthotopically engrafted with HME2-BM MT and HME2-BM shTGM2. Dotted lined indicates time point at which the primary tumor was surgically removed. (B-C) Control (GFP) and TGM2 overexpressing HME2 cells were engrafted onto the mammary fat pad via an intraductal inoculation and primary tumor growth was visualized and measured by bioluminescent at the indicated time points. Data are of individual mice resulting in the indicated P value. (D) Upon necropsy the lungs of tumor bearing mice were removed and weighed. Data in panels C and D are of individual mice resulting in the indicated P value.

**Supplementary figure 2:**
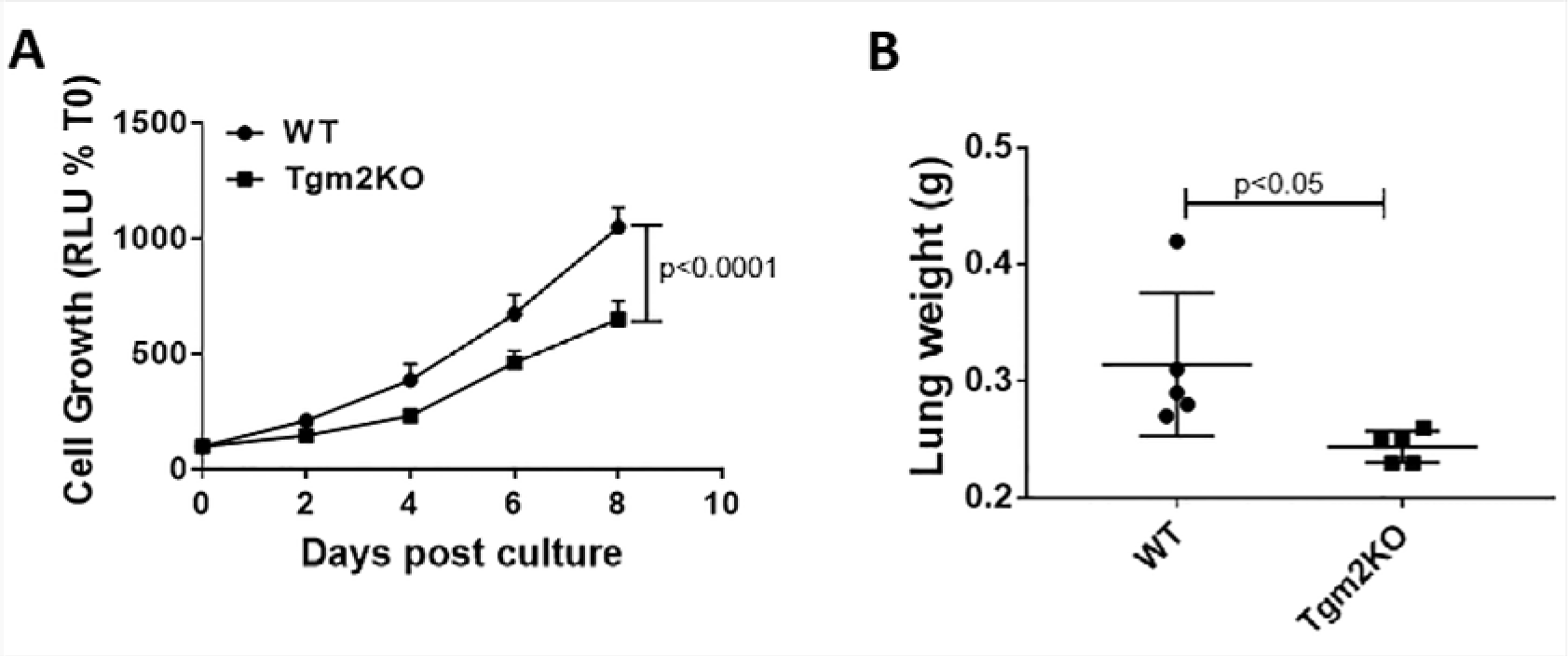
Deletion of TGM2 inhibits 3D growth and metastasis. (A) Control (WT) and TGM2 deleted (Tgm2KO) 4T1 cells were grown under single cell 3D culture conditions. Longitudinal cellular outgrowth was quantified by bioluminescence at the indicated time points. Data are normalized to the plated values and are the mean ±SD of three independent analyses resulting in the indicated P-value. (B) Control (WT) and Tgm2 deleted (Tgm2KO) 4T1 cells were engrafted onto the mammary fat pad and lungs were removed and weighed after mice were sacrificed at Day 34. Data are of individual mice resulting in the indicated mean, +SE and P value.

**Supplementary figure 3:**
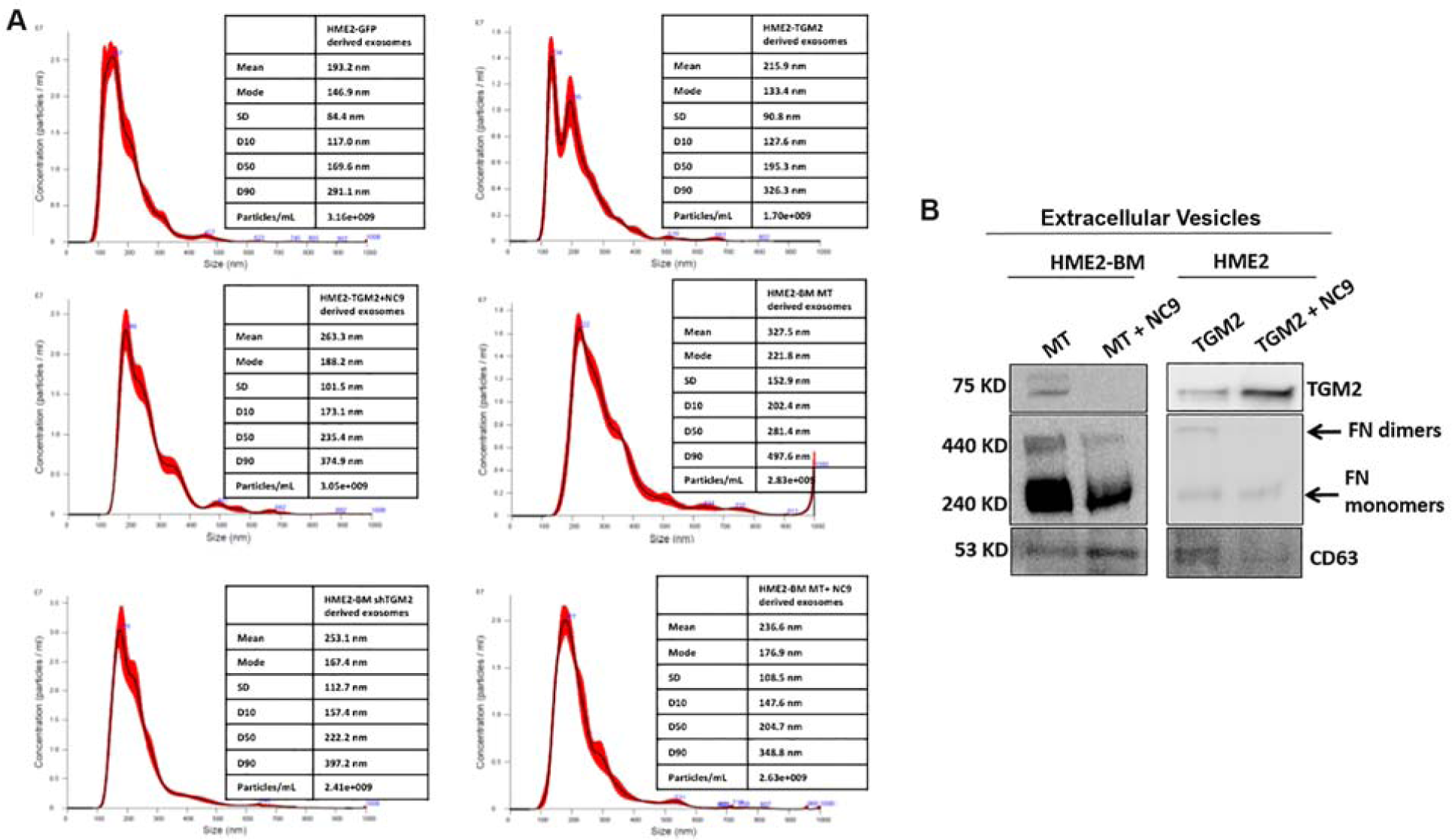
TGM2 crosslinks FN in tumor cell-derived EVs. (A) Nanoparticle tracking analysis of EVs derived from the indicated cells. (B) Immunoblot analysis of EVs derived from HME2-BM MT, HME2-BM MT treated with NC9, HME2-TGM2, and HME2-TGM2 treated with NC9. These lysates were assessed for the presence of TGM2 and FN dimerization. CD63 served as a loading control.

**Supplementary figure 4:**
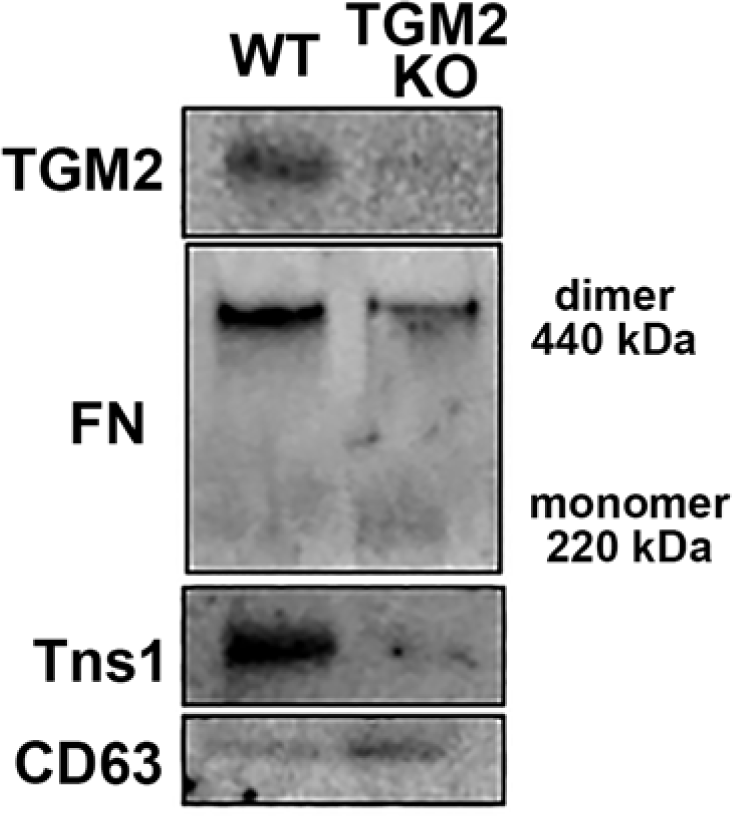
TGM2 is required for the presence of Tns1 on EVs. Extracellular vesicles were isolated from control (WT) and TGM2 deleted (TGM2KO) 4T1 cells. These EVs were analyzed for the presence of TGM2, FN dimerization, and Tns1. CD63 served as a positive control.

## Notes

Disclosure of Potential Conflicts of interest: The authors declare no conflicts of interest.

